# Resting state functional coupling between the ascending synchronising system, limbic system and the default mode network via theta oscillations

**DOI:** 10.1101/086058

**Authors:** Parnesh Raniga, Bryan Paton, Gary F. Egan

**Affiliations:** Monash Biomedical Imaging, Monash University, Melbourne, VIC 3800, Australia; The Australian eHealth Research Centre, CSIRO Health and Biosecurity, Herston, QLD 4029, Australia; School of Psychology, University of Newcastle, Callaghan, NSW 2308, Australia; ARC Centre of Excellence for Integrative Brain Function, Monash University, Melbourne, VIC 3800, Australia

**Author notes:** Equal First Authors (Parnesh Raniga, Bryan Paton). Full Address of corresponding author: Parnesh Raniga, Level 5 UQ Health Sciences Building, Royal Brisbane and Women’s Hospital, Herston, Queensland 4029 Australia.

**Keywords:** Theta oscillations, default mode network, ascending reticular activating system, Alzheimer’s disease, basal forebrain, episodic memory, autobiographical memory, memory

## Abstract

In order to better understand dysfunction in dementia and psychiatric illnesses, the underlying neuronal systems that give rise to normal memory and cognitive processes need to be better understood. Based on electrophysiological recordings in animals, theta oscillations have been proposed as an intrinsic mechanism for the orchestration of memory functions, especially episodic and autobiographical memory. Theta oscillations are controlled by the ascending synchronising system, a set of nucleui in the pontine tegmentum and basal forebrain. At a network level, the default mode network has been shown to be responsible for episodic and autobiographical.

Using resting state fMRI data, we show using an ICA approach, seed based connectivity and dynamic causal modelling that the ascending synchronising system is coupled to the medial temporal lobe nodes including the hippocampus and parahippocampal gyrus and with the default mode network. Our results provide thus support the role of theta oscillations in memory function and coordination at a network level.

**Highlights:**

- Resting state functional coupling between the DMN, MTL and ascending synchronising system.
- Theta oscillations may be the basis of this coupling given the role of these structures in control of theta.
- Theta oscillations have been implicated in memory, cognition and predictive coding.
- DMN, MTL and ASS are implicated in Alzheimer’s disease.

**Abbreviations:**

MRI: Magnetic resonance imaging
fMRI: functional magnetic resonance imaging
rs-fmri: resting state functional magnetic resonance imaging
PnO: Pontine nucleus oralis
SuM: Supra-mamillary nucleus.
MS: Medial septum
DB: Diagonal band of Broca.
VTA: ventral tegmental area
PCC: Posterior cingulate cortex
HC: Hippocampus
ARAS: Ascending reticular activating system
ASS: Ascending synchronising system
DMN: Default Mode Network
aMPFC: Anterior Medial Prefrontal Cortex
pIPL: Posterior inferior parietal lobule
NBM: Nucleus Basalis Mynert
DCM: Dynamic causal modelling
PHG: Parahippocampal Gyrus

## 1 Introduction

The neuronal systems underlying memory and related cognitive processes are of great interest not only to further our understanding of how the brain works but also to better understand dysfunction in disease states such as dementia and psychiatric illnesses. In-vivo experiments using animal models of memory formation and retrieval, and recent experiments in humans, have jointly pointed to brain oscillations in the theta (Φ) frequency ranges (4-10 Hz in mice and 1-4 Hz in humans (Jacobs, 2014)) may have a role in the synchronisation of distributed brain regions as a core mechanism of memory. These putative oscillatory mechanisms are the most likely candidate for the encoding and retrieval of short-term (Vertes, 2005), especially episodic (Burke et al., 2014; Heusser et al., 2016), and autobiographical (Fuentemilla et al., 2014) memories.

While theta rhythms have mainly been associated with the hippocampi (HC), the same rhythms can be observed in different regions of the cortex and are thought to be responsible for orchestrating long range interactions for memory formation and retrieval (Anderson et al., 2010; Fuentemilla et al., 2014; Mitchell et al., 2008). In particular, the prefrontal cortex and the posterior cingulate cortex have been observed to be entrained to hippocampal theta oscillation in humans (Anderson et al., 2010; Fuentemilla et al., 2014; Kaplan et al., 2014), primates (Tsujimoto et al., 2006) and mice (O’Neill et al., 2013) suggesting that theta entrainment may be a fundamental neuronal mechanism in mammalian brains.

At a global network level, the default mode network (DMN), a set of brain regions that are functionally connected when the brain is not engaged in an externally oriented task, has been heavily implicated in episodic memory (see (Jeong et al., 2015) for a recent review), autobiographical recall (Lin et al., 2016) and internal mentation (Buckner, 2013). Indeed, resting state functional magnetic resonance imaging (rs-fMRI) studies have shown recruitment of the hippocampus, entorhinal cortex and medial temporal lobe structures into the DMN during memory retrieval (Andrews-Hanna et al., 2010; Huijbers et al., 2011; Ward et al., 2014). Reports in literature of overlaps between brain regions displaying theta oscillations and the nodes of the DMN (Backus et al., 2016; Foster et al., 2013; Fuentemilla et al., 2014; Kaplan et al., 2014) therefore raise the question of whether a more fundamental mechanism ties the two (Foster and Parvizi, 2012).

Although theta oscillations are widely distributed in the brain, they have been shown to arise from circuits in the ascending synchronising system (ASS) (Denham and Borisyuk, 2000) in animal models. Tonic activity in the oral pontine reticular nucleus (PnO) is converted to oscillatory activity in the supramamillary nucleus (SuM). The SuM innervates the hippocampus directly and via the medial septum/ diagonal band of Broca complex (MSDB) (Borhegyi et al., 1998). Cholinergic and GABAergic cells from the medial septum show theta activity (Lee et al., 2005) and also regulate hippocampal theta (Hangya et al., 2009; Teles-Grilo Ruivo and Mellor, 2013). Moreover, dopaminergic innervation from the ventral tegmental area (VTA) to the medial septum modulates the firing and bursting rates of MSDB neurons, which in turn affect hippocampal theta activity (Fujisawa and Buzsáki, 2011; Orzeł-Gryglewska et al., 2015; Werlen and Jones, 2015). An overview of this network, thought to elicit and modulate theta activity in the hippocampus is provided in Figure 1.

**Figure 1.**
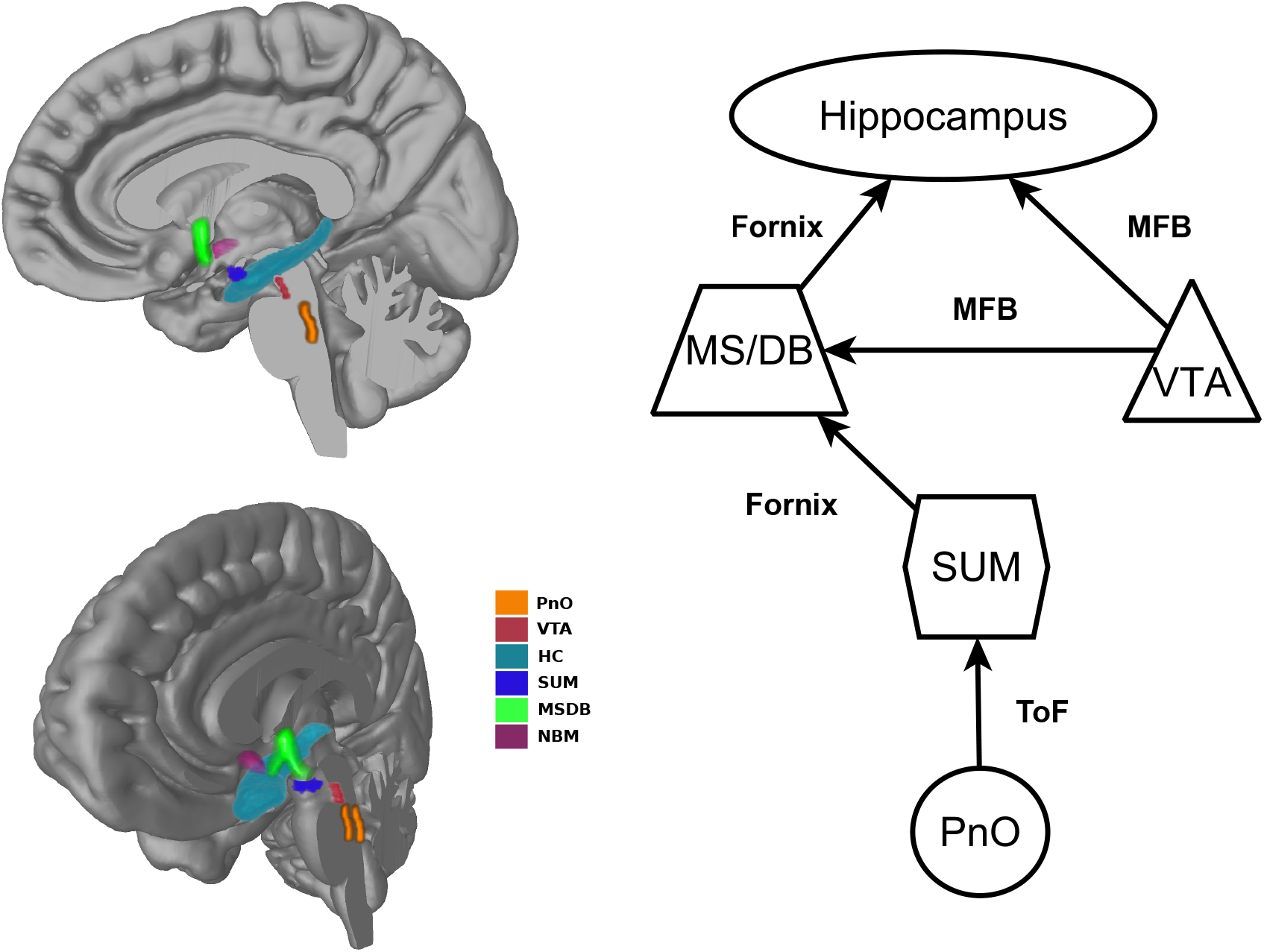
Simplified schematic representation of the parts of the ascending synchronising system implicated in the generation and maintenance of the theta rhythm (right panel) and 3D rendering of those structures in a canonical brain (left panels). Known fiber bundles connecting the regions are labelled on the schematic. These include the tract of Forel (ToF), the fornix and the medial forebrain bundle (MFB).

Theta oscillations are thought to be one of the core mechanisms in memory processes (Düzel et al., 2010) including encoding, retrieval and autobiographical introspection. Understanding the functional role of theta oscillations may enhance our understanding of normal brain function and dysfunction in disease states such as Alzheimer’s disease. The pathologies of AD including atrophy in the medial temporal lobe (Grothe et al., 2016), the basal forebrain (atrophy (Kerbler et al., 2015), tau pathology (Schöll et al., 2016), and loss of cholinergic function (Mufson et al., 2008)) and beta-amyloid build-up in the DMN (Buckner et al., 2005) exhibit anatomical patterns that overlap with nodes showing theta oscillations. Moreover, the earliest known neurofibrillary tangles are observed in the ascending reticular activating of which the ascending synchronising system is a part (Braak et al., 2011).

While theta generation and modulation networks have been extensively studied in animals, they are more difficult to study in humans due to the difficulty of in-vivo electrophysiological measurements and the widespread nature of the network. Electrophysiological recordings, due to their inherent local nature, have been limited in mapping the theta activity at the network and global levels. Macroscopic approaches such as EEG and MEG have shown the involvement of the DMN but do not have the spatial resolution to resolve the synchronising network. We investigated whether these synchronisation networks could be observed using high temporal and spatial resolution resting state fMRI and if they were coupled to the default mode via limbic nodes that have previously been shown to be coupled to the DMN. Identifying these networks could provide evidence for the role of theta oscillations in orchestrating dispersed cortical regions into extended brain networks supporting memory function.

Using resting state data from 300 subjects from the Human Connectome Project (Van Essen et al., 2013), we show using a group PCA-ICA approach that the PnO, VTA and MSDB can be observed to be coupled with MTL structures and the DMN. Further to this, we show with functional connectivity using seeds from the PnO, SuM, VTA and MS, functional networks that are consistent with the DMN. We then use a dynamic causal modelling (DCM) analytical approach to show that the DMN incorporating the HC, MSDB, SuM, PnO and VTA is the most likely network given the observed data. To our knowledge, this is the first report of the coupling between the ASS including MSDB, MTL and DMN in humans using rs-fMRI and provides evidence that theta oscillations may serve as a brain mechanism for coordinating these disparate brain regions.

## 2 Results

### 2.1 Group PCA-ICA

A default mode network ICA component was isolated from the group based PCA-ICA results based on visual inspection. The network for Band 1 (0.001-0.027 Hz) (see Figure 2A and Supplementary Figure 1) showed a pattern consistent with the DMN with functional connectivity observed between the posterior cingulate cortex (PCC), anterior middle prefrontal cortex (aMPFC), and posterior inferior parietal lobule (pIPL). Furthermore, limbic structures including the bilateral hippocampii (HC), the retrospinal cortex (RSP) and parahippocampal gyrus (PHG) were also observed. The medial septum region was also functionally connected to the DMN along with the nucleus basalis of Meynert (NBM), the latter only being connected in Band 1.

**Figure 2.**
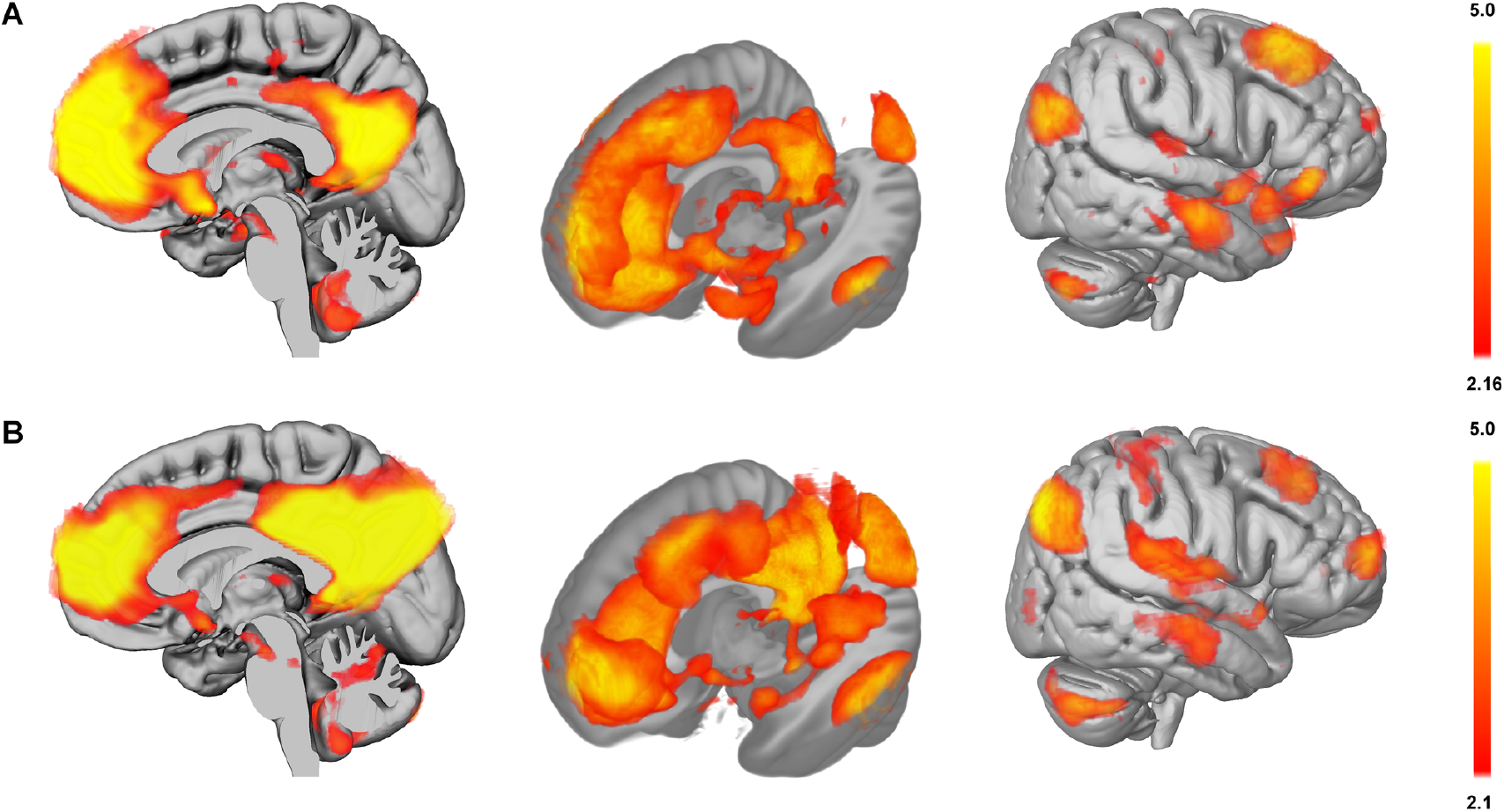
The default mode network as observed in Band 1 (0.01-0.027 Hz) (A) and Band 2 (0.027-0.07 Hz) (B). Overlays are thresholded z maps for the components as produced by melodic from FSL. In Band 1, the medial septum/diagonal band of broca complex, ventral tegmental area and midline thalamus are connected to the default mode and the hippocampus. In Band 2, the medial septum/diagonal band of broca complex, ventral tegmental area, pontine nucleus oralis and midline thalamus are connected to the default mode and the medial temporal lobe.

The network for Band 2 (0.027-0.07 Hz) (Figure 2B and Supplementary Figure 1) showed similar patterns of functional connectivity as for Band 1. In the mid brain and pontine tegmentum, two distinct but connected regions of functional connectivity could be observed. The rostral region was identified to be the ventral tegmental area, which is consistent with literature reports of functional connectivity between the VTA and the DMN (Fujisawa and Buzsáki, 2011; Werlen and Jones, 2015) in animal models and in fMRI (Tomasi and Volkow, 2014; Tompary et al., 2015). The VTA was also evident in Band 1. The adjacent activation region was more caudal and dorsal, consistent with the location of the rostral portion of the pontis nucleus oralis (PnO) (see Figure 3 for comparison with location of the PnO from the Harvard Ascending Arousal Network Atlas (Edlow et al., 2012)). Functional connectivity with only the rostral portion of the PnO is consistent with the literature on animal stimulation for the elicitation of hippocampal theta (Vertes and Kocsis, 1997).

**Figure 3.**
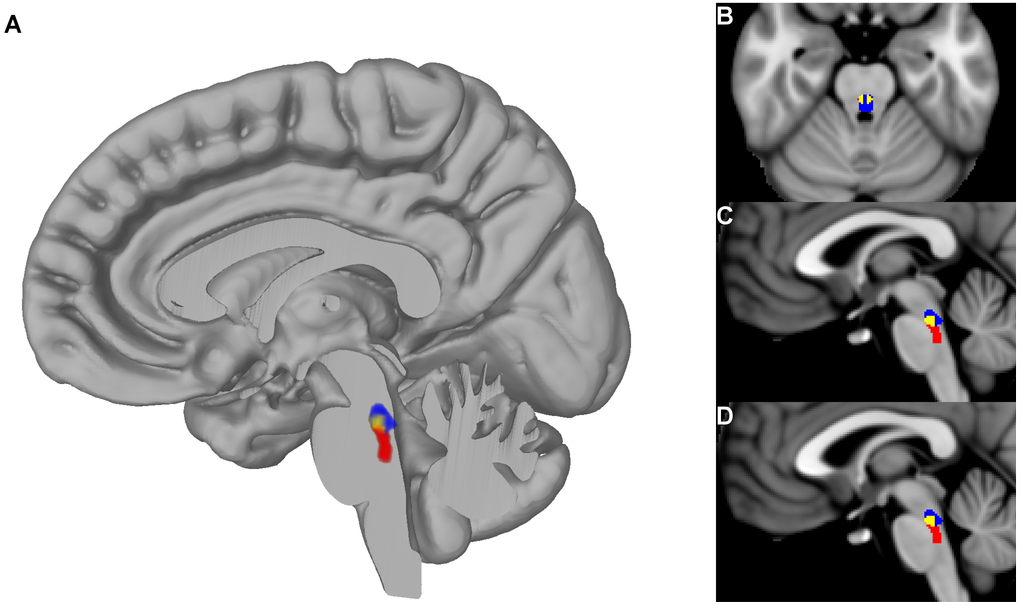
PnO ROI as defined from the group PCA-ICA (shown in blue), ROI from the Harvard Ascending Arousal Network (AAN) atlas (shown in red) and overlap in yellow shown in a 3D rendering (A), axial section B (MNI coordinates Z=-24), sagittal sections C (MNI coordinates X=−1) and D (MNI coordinates X=2).

The mammillary bodies were functionally connected in both Band 1 and 2. Also, the DMN in both Bands 1 and 2 had prominent activations of the midline thalamus with activations in the Band 1 DMN being along the entire midline of the thalamus, but restricted to a region just posterior of the thalamic adhesion in the Band 2 DMN. The DMN results for Bands 3, 4 and 5 are not reported here, as the typical resting state correlational analysis is limited to 0.01 to 0.1 Hz. Apart from the DMN structures themselves, no other structures were observed to be coupled to the DMN at other frequencies.

To cross-validate the results we performed global seed based analysis on a different set of 100 subjects using seeds based on literature and the PCA-ICA results (see Table 2), and a frequency band typical of resting state data (0.01-0.1Hz).

**Table 1.**
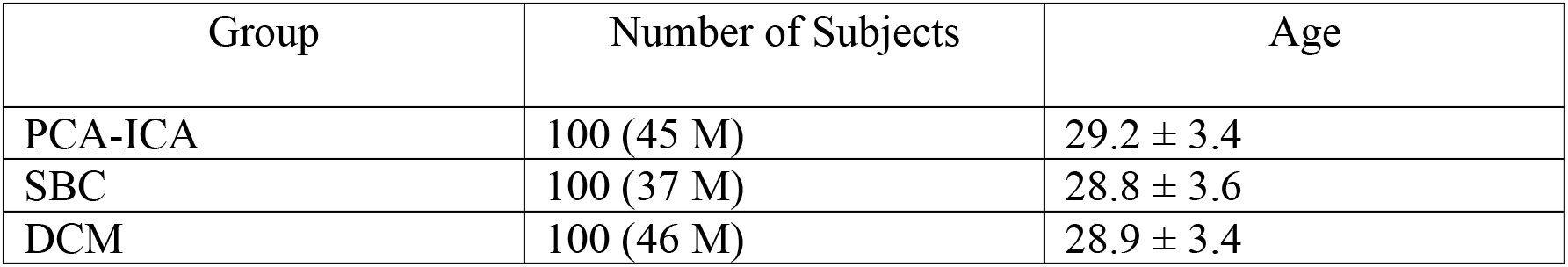
Demographic information of the HCP subjects used in the study. Age of subjects is taken at the centre of the age range of the subject from the Human Connectome Project. For subjects in the over thirty-six years age range, the age was taken to be thirty-six years.

**Table 2.**
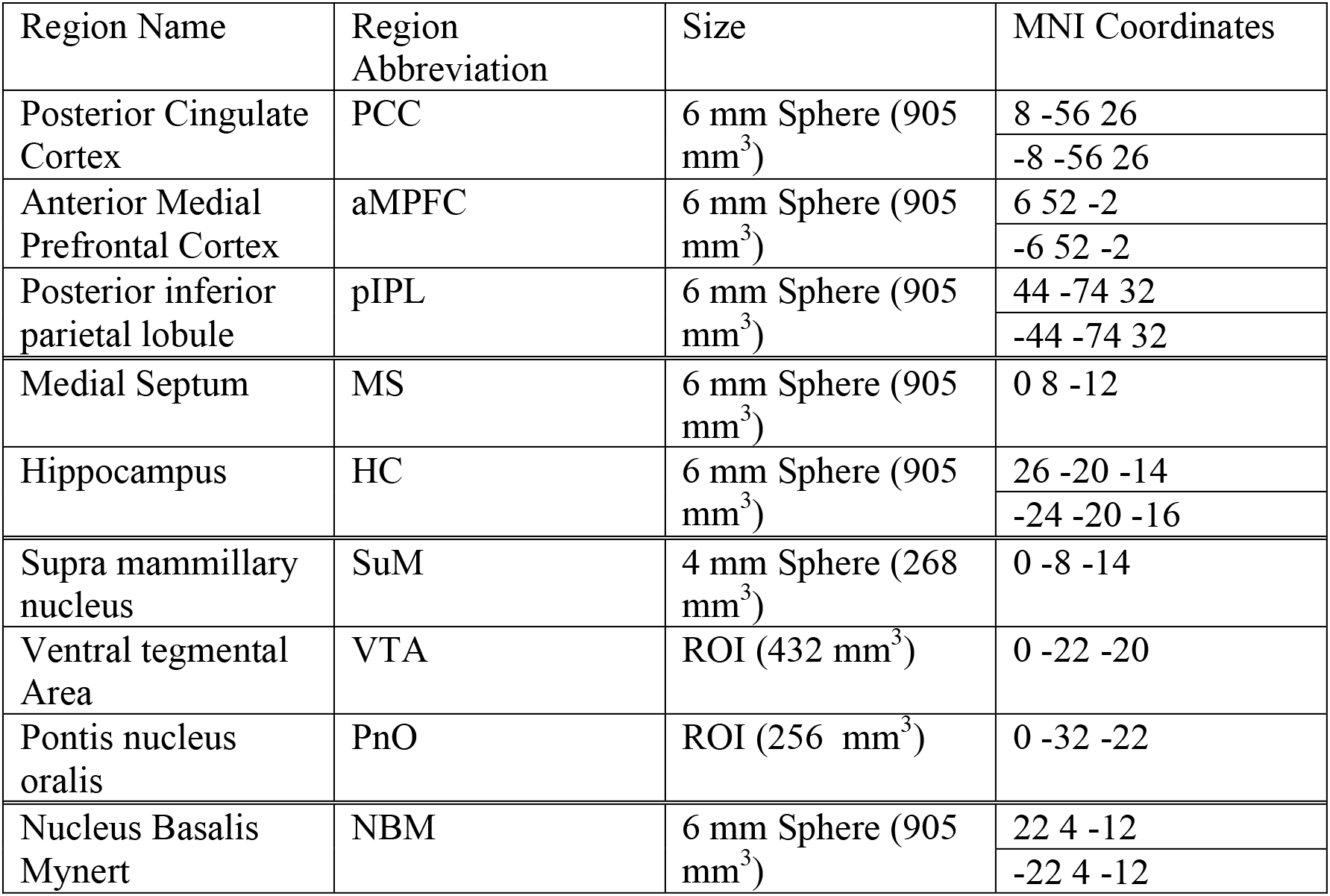
Regions of interests used in the seed based connectivity and dynamic causal modelling, their sizes and MNI coordinates. Where bilateral ROIs were used, both left and right coordinates are provided. Spheres with a 6mm radius were used for all regions except the supra mammillary nucleus where a 4mm sphere was used to avoid signal spill over and the VTA and PnO where the ROIs were voxel masks extracted from the PCA-ICA analysis.

### 2.2 Seed based Connectivity

Voxel based connectivity patterns for seed regions from the MSDB, PnO, SuM and VTA are presented in Figure 4 (and also Supplementary Figure 2). Connectivity for MSDB, PnO, SuM and VTA showed a profile consistent with regions composed of the default mode, whilst the MSDB and VTA showed strong connectivity to DMN regions as well as limbic regions including hippocampus and retrospinal cortex.

**Figure 4.**
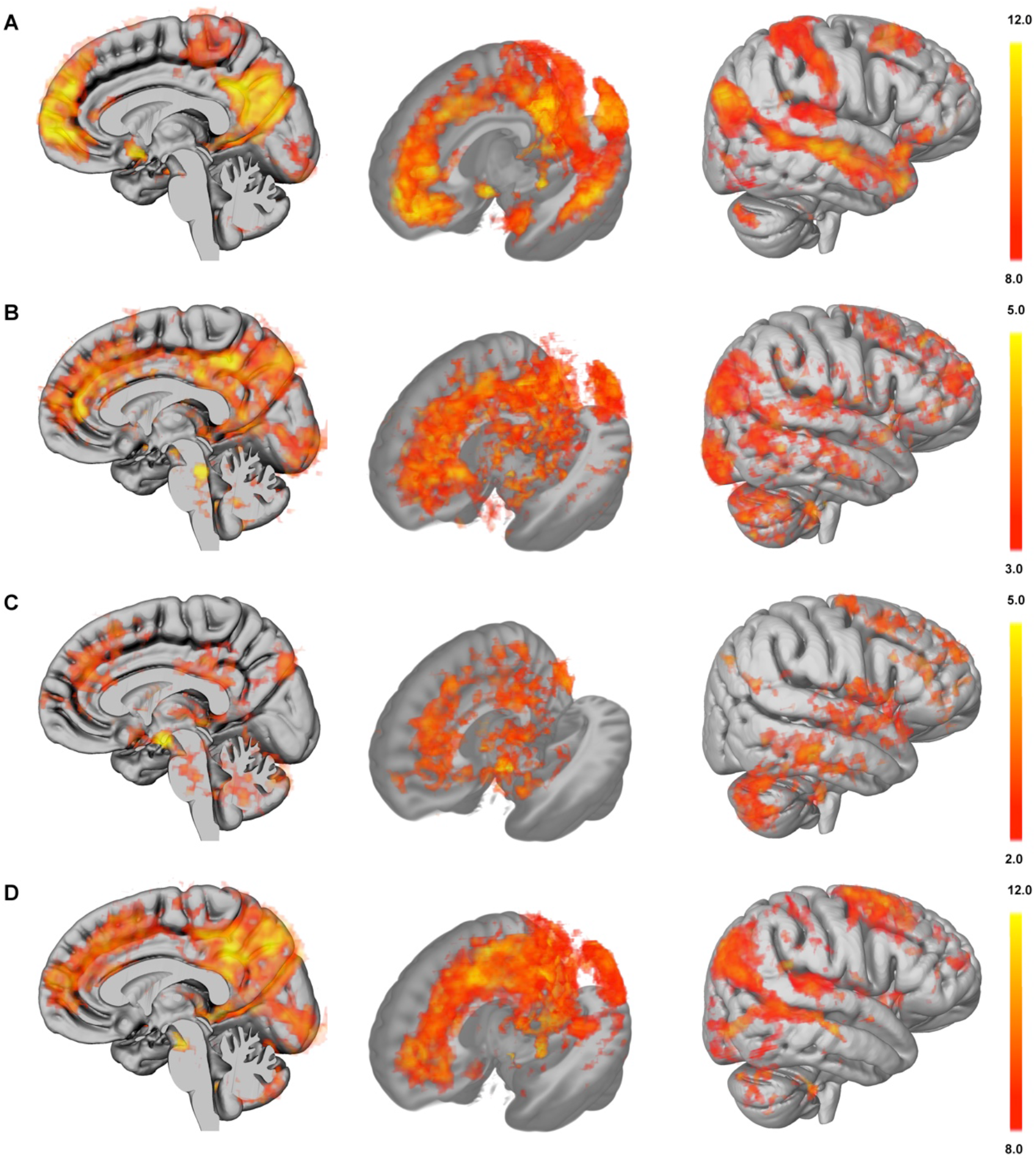
Seed based connectivity. T-scores are displayed for (A) MSDB, (B) PnO, (C) SuM and (D) VTA. All voxels displayed are significant at p < 0.05 using fsl-randomise permutation tests and threshold free cluster enhancement, however higher thresholds are used in the display to show relevant details.

The PnO showed a pattern of connectivity consistent with the DMN although the activation was noisier than for the VTA and MSDB. The SuM connectivity pattern, while showing regions overlapping with the DMN, was very noisy likely due to the fact that the SuM is a very small region with a highly heterogeneous signal.

The connectivity of the NBM was consistent with that of the salience network with functional connectivity observable across the cingulate sulcus and along the dorsal anterior cingulate cortex. Strong functional connectivity was also observed with bilateral insula (see Supplementary Figure 3).

Overall, the seed based functional network connectivity patterns mirror those observed using the group PCA-ICA method, providing empirical replication of the specific activity and in particular the correlated but wide spread activity in the DMN. To determine if these results were an epiphenomena we fitted and inverted a series of DCM models of increasing complexity with the aim of determining the best model that could causally explain the observed functional connectivity.

### 2.3 Dynamic Causal Modelling

Although the functional connectivity measures are indicative of coupling between the DMN, limbic nodes and the ASS, fitting a physiologically informed causal model of the dynamic patterns of connectivity may be able to infer the true network topology and the underlying physiological parameters. We fitted five different models (see Figure 5) that included the DMN, limbic regions and the nodes of the ASS and inverted the models using spectral DCM (Friston et al., 2014) in a group of 100 subjects. The models were compared using Bayesian Model Selection (Rigoux et al., 2014).

**Figure 5.**
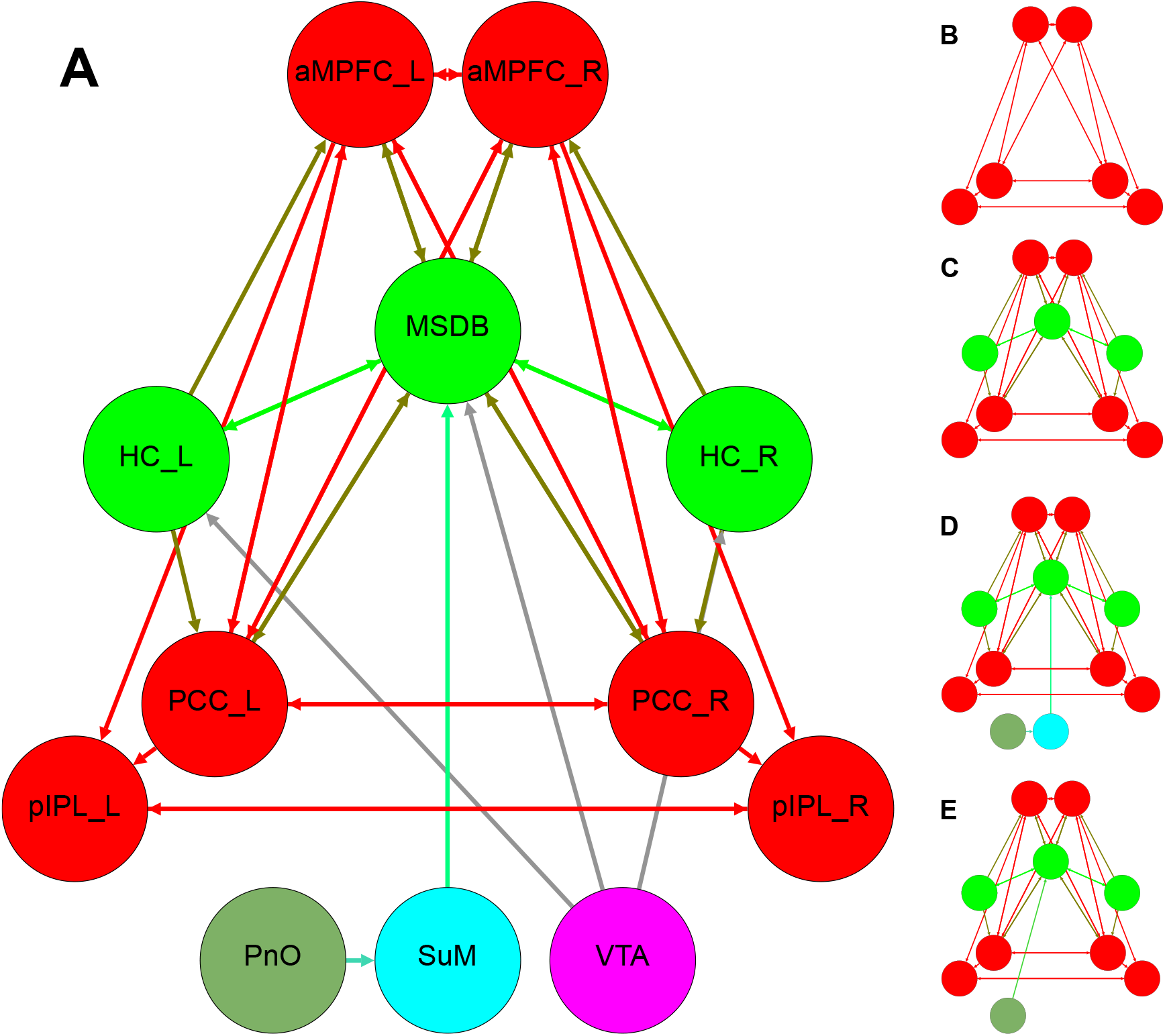
The models tested as part of the DCM analyses. Nodes belonging to the classical default mode are shown in red, those belonging to the limbic system are shown in green, the PnO is shown in olive, the SuM is shown in turquoise and the VTA in purple. The models are of increasingly greater complexity with Model 1 (B) including only the classic default mode (red nodes), Model 2 (C) including the DMN and limbic nodes (red + bright green nodes) and Model 3 (D) including the DMN, limbic structures and ascending synchronising system sans the VTA (red, bright green, olive and turquoise nodes). Model 4 (E) proposes a connection between the PnO and the MSDB (red, bright green, olive nodes) and Model 5 (A) includes the VTA as connected to the limbic structures.

The basic DCM model, model 1, included the cortical nodes of the DMN (Sharaev et al., 2016). Model 2 increased the complexity of the DMN by inclusion of bilateral hippocampi and the MSDB with reciprocal connections to all nodes. Model 3 included the nodes of the ascending synchronising system connected to the MSDB. Model 4 removed the SuM as that region had the least consistent seed based connectivity pattern. Model 5 was the most complex with the VTA connected to the MSDB and the hippocampus based on anatomical connectivity in animal models.

Results of the DCM analysis showed that the log evidence of the five models were M1=2.5159e6, M2=2.5579e6, M3=2.5597e6, M4=2.5579e6 and M5=2.5795e6. Further, the results showed that the most likely model given the data from those models tested is Model 5 (model posterior probability of 1.0). Model 5 included the HC, MSDB and the theta generation nodes of the ASS (PnO, SuM and VTA) connected to the DMN, could explain 90±4% of the variance in the data (see Supplementary Figure 4 for boxplots of variance explained for all models). Also Model 3 had higher log evidence and explained greater variance compared to Model 4 suggesting that the SuM may play a role in this coupled network.

## 3 Discussion

Our results provide the first in-vivo evidence of functional coupling between the ascending synchronising system, limbic regions including the HC, PHG, MSDB and the classical DMN. While the integration of MTL structures including HC and PHG into the DMN has already been observed (Andrews-Hanna et al., 2010; Ward et al., 2014) and the inclusion of the MSDB has been described (Greicius et al., 2003), the coupling of the ASS together with the MTL and DMN in humans identified using rs-fMRI is novel.

The results of the frequency specific group PCA-ICA showed typical patterns for DMN connectivity across all frequency bands with the PnO observable only in Band 2 (0.027-0.7 Hz) and the NBM observable only in Band 1 (0.01-0.027 Hz). The MSDB and VTA were prominent in both Bands 1 and 2. To cross-validate the coupling we derived and to ensure that the coupling we saw was not an artefact of the group PCA-ICA method (lower variance compared to standard ICA approaches) or of the parameters we chose (low-dimensionality), we performed seed based, whole brain connectivity analysis in a different subset of subjects. Functional connectivity from the PnO, VTA and MSDB showed statistically significant patterns of connectivity typical of the DMN. While the SuM showed similar but noisier patterns, clusters concomitant with the pIPL were not present in the SuM results. It is possible, due to the size of the SuM and the proximity of other nuclei that that the sampled signal was too heterogeneous. However previous studies have observed a strong resting state functional connectivity between the mammillary bodies and the HC (Blessing et al., 2016).

Further cross-validation using DCM revealed that of the five models tested, the most likely model given the data was one where the nodes the of ASS (including SuM, PnO and VTA) and the limbic system (HC and MSDB) were coupled. The Bayesian procedures used in the DCM model fitting balance the relative accuracy of a model with the complexity of the same model, a form of Occam’s razor. A model that has many parameters can more closely fit the data but has poor generalisability, whereas a more parsimonious model might fit the data slightly worse but can be readily applied across many situations. In line with this accuracy versus complexity balance, the inclusion of SuM resulted in a better model fit and greater explained variance even when including the extra parameters (Model 3 vs. Model 4). This highlights the causal role of the SuM in supporting the memory processes through theta based mechanisms as has been suggested from animal studies.

While the hippocampus and medial temporal regions such as the entorhinal cortex, parahippocampal regions are known to be the core regions for the retrieval of episodic and working memory (Ritchey et al., 2015) and have been shown to be coupled to the DMN (Andrews-Hanna et al., 2010; Ward et al., 2014), the neural mechanisms underlying memory processes and coupling are not well understood. Oscillations in neuronal firing over the theta frequency band, 4-10 Hz in mice and 1-4 Hz in humans (Jacobs, 2014) have been proposed as a candidate mechanism for not only memory function but long range communications in memory related neural circuits (Gollo et al., 2011). In particular, theta oscillations have been shown to be relevant for memory, both episodic and short term (Burke et al., 2014; Fuentemilla et al., 2014; Heusser et al., 2016; Vertes, 2005). Not only are the distribution of the theta rhythms important, but memory performance tracks the power of theta oscillations (Backus et al., 2016), and coupling between the MTL and medial prefrontal cortex, in the theta range, has been shown to be present in memory encoding and retrieval (Anderson et al., 2010; Backus et al., 2016; Fuentemilla et al., 2014; Kaplan et al., 2014; Lin et al., 2016).

Whether theta oscillations are generated from a set of pacemaker neurons or are intrinsic properties of the coupling of certain neuron populations (Buzsáki, 2002) is unclear. Experiments on animal models have shown that theta power (Orzeł-Gryglewska et al., 2015; Vertes and Kocsis, 1997; Werlen and Jones, 2015) and subsequent memory performance appear to be modulated by a set of nuclei including the PnO, SuM, VTA and MSDB, termed the ascending synchronising system (Denham and Borisyuk, 2000). The functional coupling of these structures to the DMN in rs-fMRI supports the hypothesis that theta oscillations are one of the underlying mechanisms for episodic and autobiographical memory (Burke et al., 2014; Buzsáki and Moser, 2013; Fuentemilla et al., 2014) and potentially the neurobiological substrate for long range coordination of memory interactions (Gollo et al., 2011).

The findings also support the role of theta rhythms to contextualise prediction error processing in medial temporal lobe structures coupled to cortical nodes of the DMN via theta oscillations (Carhart-Harris and Friston, 2010; Carhart-Harris et al., 2014; Friston, 2010). This coupling allows for the integration of primary sensory information with memory and contextual information for higher order processing and thus may form a substrate for the waking adult conscious experience. The slower theta oscillations provide a context in which prediction error units can signal (via gamma band bursts) discrepancies between predicted and actual sensory inputs. The roles of this coupling is critical and its breakdown may underlie unusual and altered states of consciousness, as seen in psychosis for example (Carhart-Harris and Friston, 2010; Carhart-Harris et al., 2014).

Our observations about brainstem coupling to the DMN are however not new, with two recent papers on intrinsic hippocampal functional connectivity (Bär et al., 2016; Blessing et al., 2016) showing similar patterns of functional connectivity with medial temporal lobe structures and also to the DMN. In particular, both these studies, in different populations using different acquisition and processing paradigms, have shown functional connectivity of the VTA, MSDB (labelled the subcallosal area) to a region in the brainstem labelled as the median raphe nucleus (MRN). The raphe nuclei are the source of serotonin release in the brain with projections throughout the entire brain (Wagner et al., 2016). Given that activation of the MRN tends to desynchronise hippocampal theta activity and serotonin depletion tends to increase theta EEG power and working memory function (López-Vázquez et al., 2014), we suspect that the relevant brainstem region may be the PnO. Moreover, since in humans the PnO surrounds the MRN the two regions can be easily misidentified due to partial volume effects and motion at the resolution of typical functional MRI experiments (see Supplementary Figure 5).

While nuclei such as the ventral tegmental nucleus of Gudden (VTNg) have been thought to be a part of the limbic system and theta synchronising system (Kocsis et al., 2001), the VTA as a node in this system has been recently proposed (Orzeł-Gryglewska et al., 2015). In particular, the VTA is thought to help synchronise cortical and subcortical interactions (Fujisawa and Buzsáki, 2011). In light of these findings and our results, the VTA and dopamine may play a role in novelty detection and long term potentiation for memory encoding (Lisman and Grace, 2005) and working memory (Fujisawa and Buzsáki, 2011). Moreover, with evidence of phase locking, where bursts of activity between two regions are synchronised, between the VTA and hippocampal theta (Fujisawa and Buzsáki, 2011; Orzeł-Gryglewska et al., 2015), the activation of the VTA with the DMN is consistent with our model. The coupling between the hippocampus and the VTA has recently been shown to provide context to rewarding experiences (Luo et al., 2011). Moreover, increased connectivity between VTA and MTL regions has been linked to memory consolidation (Tompary et al., 2015).

How can these rs-fMRI findings be interpreted? It is unlikely that the temporal resolution and the slow response of the hemodynamic system would enable direct observations of theta oscillations using rs-fMRI. However, unlike in mice and rats, the power of theta oscillations in humans is not continuous but rather occurs in more discrete bursts lasting about a second (Watrous et al., 2013). Hemodynamic coupling of such bursts, reflecting envelope changes in theta oscillations would result in detectable BOLD changes at around 0.1 Hz (Foster and Parvizi, 2012). This frequency of BOLD fluctuations is at the upper edge of the usable spectrum of typical fMRI studies (assuming a 2 s sample rate [TR]) but well within the spectrum of the data used here. The coupling of the VTA and its hypothesised role in producing bursts of theta activity also supports this notion (Fitch et al., 2006; Fujisawa and Buzsáki, 2011). Furthermore, the BOLD contrast has been shown to be most directly correlated with power of high frequency local field potentials (Logothetis et al., 2001). Since theta and gamma band oscillations are known to be coupled (Lisman and Jensen, 2013) and recent evidence shows this coupling is active in episodic sequence memory (Heusser et al., 2016), it is likely that coupled gamma band oscillations would give rise to the BOLD response changes in the low frequency ranges.

While this network has been observed previously, the number of subjects, length of rs-fMRI acquisitions, quality of the data and the state of the art pre-processing method have combined to produce the current results. The rs-fMRI acquisitions were acquired for a duration of 15 mins per run which is not only longer than typical acquisitions but also has higher temporal signal to noise due to the use of multi-band acquisition protocol. Furthermore, the acquisitions were phase encoded left to right and not anterior to posterior. This phase encoding scheme reduces the signal pileups and dropouts in regions close to the air tissue interfaces such as the MSDB, the VTA and mammillary bodies. Coupled with this, the standardised pre-processing pipeline utilised in the HCP, along with a flip angle close to the Ernst angle, ensures that confounds such as motion and cardiac and respiratory signals have a drastically reduced influence on the signal of interest.

The coupling of the ASS, MTL and the DMN via theta oscillations in resting state has implications not only for the understanding of how the brain functions normally but also of how it is affected in neurological and psychiatric disorders. Pathological and functional changes in the default mode have been observed in several disorders, most prominently in Alzheimer’s disease (Buckner et al., 2005) whereby amyloid beta plaques form in regions overlapping the default mode (Grothe et al., 2016). Moreover, neurofibrillary tangles and atrophy are more prominent in MTL structures with NFTs initially appearing in the pontine tegmentum and midbrain followed by the entorhinal and hippocampal cortices (Braak et al., 2011) and in the basal forebrain. Moreover, theta oscillations have been observed to be selectively increased in mice models of Alzheimer’s disease and in humans, EEG signal in the theta range increases in MCI and AD subjects (Hamm et al., 2015).

While in the current study, we have been able to show coupling between the default mode network, medial temporal lobe structures and the ascending synchronising system, our hypothesis that this was orchestrated by theta oscillations is speculative based on prior work and we are unable to directly measure the oscillations using fMRI. Also, while we cross-validated our PCA-ICA results using seed based global connectivity, the resultant maps were noisy compared to the PCA-ICA maps and other maps that have been produced prior although they clearly showed patterns consistent with the DMN. This may be due to the lower smoothing (FWHM 4 mm) that we employed and the size and heterogeneity of seed voxels. Nevertheless, this is a potential shortcoming that we aim to further explore in future work.

The particular form of the DCM model used, spectral DCM, is based upon the cross-spectral density between the various nodes of the fitted networks. The reasoning being that the second order statistics, the cross spectral densities are all that are needed to explain the coupling between the network nodes under steady state assumptions (Friston et al., 2014). The switch from estimating hidden neuronal states from nodal time series, where in the case of resting state fMRI data there are unknown connection weights and an unknown number of network nodes and stochastic time series fluctuations, to a problem of estimating the cross spectral densities and their associated noise transforms the estimation problem into a deterministic one (Friston et al., 2014). This switch also ensures a time and computationally efficient model estimation process, especially in the case of this current study given the modest number of networks, the density of the time series and the large number of participants. One limitation of the spectral DCM is the steady state assumption although this greatly improves the model evidence estimation process it means that short lived fluctuations and modulations of connection strengths are not identifiable.

In conclusion, we have presented in first *in-vivo* results in humans showing coupling between the ascending synchronisation system, the hippocampus and MTL structures and the typical DMN nodes using rs-fMRI. This was done using a purely data-driven approach, namely group PCA-ICA and confirmed using two different hypothesis driven approaches namely seed based connectivity and dynamic causal modelling. Our results add to the growing body of literature supporting the role of brain oscillations in the theta frequency range as being the neurobiological basis of episodic and autobiographical memory. Furthermore, we have discussed the potential role of this coupled system in AD with differing pathologies affecting different parts of the coupled system and the major impact of the disease being on memory, the system hypothetically supported by this coupled system. In future work, we aim to explore the implications of this coupling for neurological and psychological conditions.

## 4 Materials and Methods

### 4.1 Subject Information

Resting state fMRI data from 300 subjects from the human connectome project from the S500 release was used in this study. The subjects were split into three groups each of 100 subjects to cross-validate the three sets of analyses, namely: (i) the group PCA-ICA, (ii) seed based connectivity, and (iii) dynamic causal. The rs-fMRI data was processed by the minimal preprocessing pipeline as used in the Human Connectome Project (Glasser et al., 2013). A single run of the rs-fMRI dataset from each subject was used in this study. For all subjects, the first run, which was acquired with a phase encoding from left to right was used. Ethics approval for the HCP project and protocol was granted by the local Institutional Review Board at Washington University in St. Louis. Full details on the HCP data set have been published previously (Van Essen et al., 2013).

### 4.2 MRI data

The rs-fMRI data was processed by the HCP minimal preprocessing pipeline (Glasser et al., 2013). Briefly, for rs-fMRI datasets, motion correction was performed and the 12 motion parameters were regressed out of the data. Image artefacts were removed using an ICA based, machine learning technique (Griffanti et al., 2014). The rs-fMRI dataset was registered to the MNI space.

### 4.3 Band-Specific Group PCA-ICA

The datasets of each subject from the first group of one hundred subjects was bandpassed into five bands (Gohel and Biswal, 2015) [(0.01-0.027), (0.027-0.073), (0.073-0.198), (0.198-0.5), (0.5-0.69)] Hz using the 3dBandpass program from the AFNI suite (Cox, 1996). The bandpassed images were smoothed using a Gaussian smoothing function with a FWHM of 5mm using SUSAN (Smith and Brady, 1997).

For each band, images of all the subjects (100 subjects) were fed into melodic (Smith et al., 2014). A modified version of the program from FSL version 5.0.8 was used whereby multicore computations were implemented for the PCA step of the algorithm. All other steps and code remained identical. The number of extracted components was fixed at 15 components to ensure that complete networks and not partial networks were extracted.

### 4.4 Seed based connectivity

Regions of interest were created based on the work of Andrews-Hanna and coworkers (Andrews-Hanna et al., 2010) with the ICA maps of the DMN extracted from the group PCA-ICA. In all, 11 regions of the DMN and the nucleus basalis of Mynert were created bilaterally as spheres of 6 mm radius. Regions for the medial septum/vertical limb of the diagonal band of broca (MSDB), supra mammillary nucleus (SuM), pontis nucleus oralis (PnO) and the ventral tegmental area (VTA) were created with their sizes and MNI coordinates presented in Table 2.

Voxel based correlation analysis was undertaken for ROIs including the PnO, SuM, MS, VTA and NBM. For each subject, a time-course corresponding to each region was extracted using 3dmaskSVD. Following this, the rs-fMRI datasets were detrended using 3dDetrend, bandpassed using 3dBandpass to a passband of 0.01 – 0.1 Hz and smoothed using a Gaussian filter with a FWHM of 5 mm using 3dBlurToFWHM.

The extracted time-courses were correlated globally with the post-processed rs-fMRI data. Whole brain correlation coefficients were tested for significance via a one sided test using permutation testing and threshold free cluster extent (Smith and Nichols, 2009; Winkler et al., 2014). Testing was performed using the randomise tool from FSL and fifty thousand permutations were performed. These steps were also applied to the data for the second group of one hundred subjects.

### 4.5 Dynamic Causal Modelling

Time courses were extracted from the rs-fMRI volumes using the ROIs from the seed based connectivity analyses above.

The DCM models were constructed based on the following assumptions:

1. The basic default mode consisted of the PCC and aMPFC connected bilaterally and reciprocally. The pIPL regions were connected unidirectionally to the PCC and aMPFC regions ipsilaterally and connected reciprocally to each other as per the results of Sharaev and coworkers (Sharaev et al., 2016).
2. The hippocampus was connected to the PCC and aMPFC unidirectionally and hemispherically.
3. The MSDB was connected to all regions reciprocally.
4. The PnO was connected to SuM unidirectionally and the SuM was connected to the medial septum unidirectionally.
5. The VTA was connected to the MSDB and to the hippocampus unidirectionally.

Based on these assumptions, five anatomical network models were constructed. Model 1 was the base model of the DMN (assumption 1). Model 2 included the hippocampus and medial septum (assumptions 1, 2, 3). Model 3 included the theta generation aspect of the ascending synchronising system but the VTA was not connected (assumptions 1,2,3,4). Model 4 was the same as Model 3 but with direct connectivity between the PnO and MS thereby excluding the SuM. Model 5 was the same as Model 3 but included the links between the VTA and MSDB and hippocampus. The five models are shown in Figure 5.

The models were inverted using the spectral DCM method (Friston et al., 2014) and compared to each other based on Bayesian model selection as well as in terms of explained variance.

## 5 Acknowledgments

Data were provided by the Human Connectome Project, WU-Minn Consortium (Principal Investigators: David Van Essen and Kamil Ugurbil; 1U54MH091657) funded by the 16 NIH Institutes and Centers that support the NIH Blueprint for Neuroscience Research; and by the McDonnell Center for Systems Neuroscience at Washington University.

We would like to acknowledge the input of Francesco Sforazzini from Monash Biomedical Imaging into some aspects of the analysis of the imaging data.

---

**Supplementary Figure 1.**
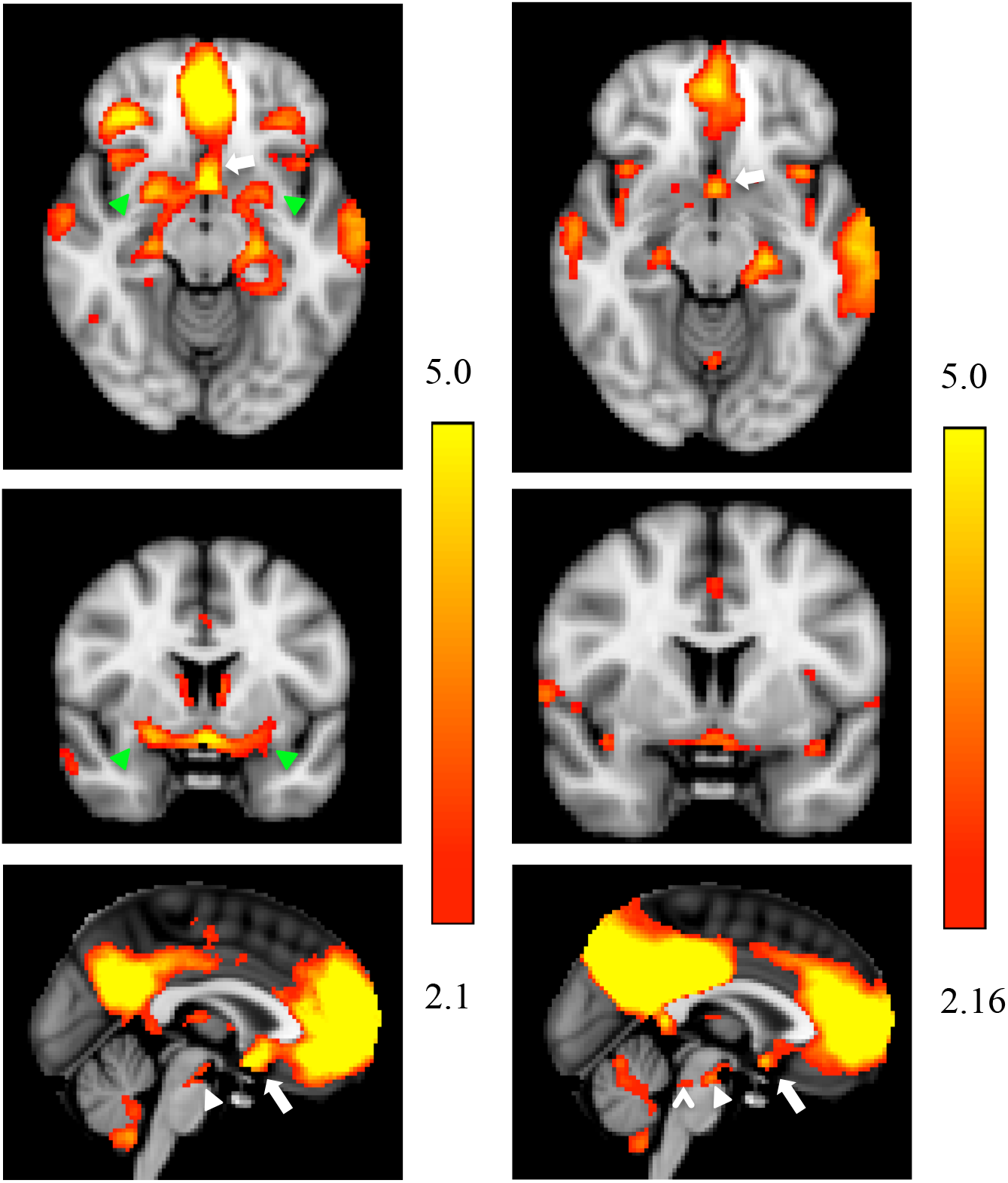
Cortical and subcortical structures that are a part of the default mode at Band 1 (0.001-0.027 Hz) left column and Band 2 (0.027-0.073 Hz) right column. Axial slices (z= −12mm), coronal slices (y=4mm) and sagittal slices (x= 0 mm) in the MNI coordinate system illustrate known cortical regions including the hippocampus and the para-hippocampus cortex and medial regions in the thalamus but also ventral tagmental area in the midbrain (white arrowhead), the medial septum in the forebrain (white arrow), the nucleus basalis of mynert (NBM) (green arrowheads) and the pontine nucleus oralis (carat)

**Supplementary Figure 2.**
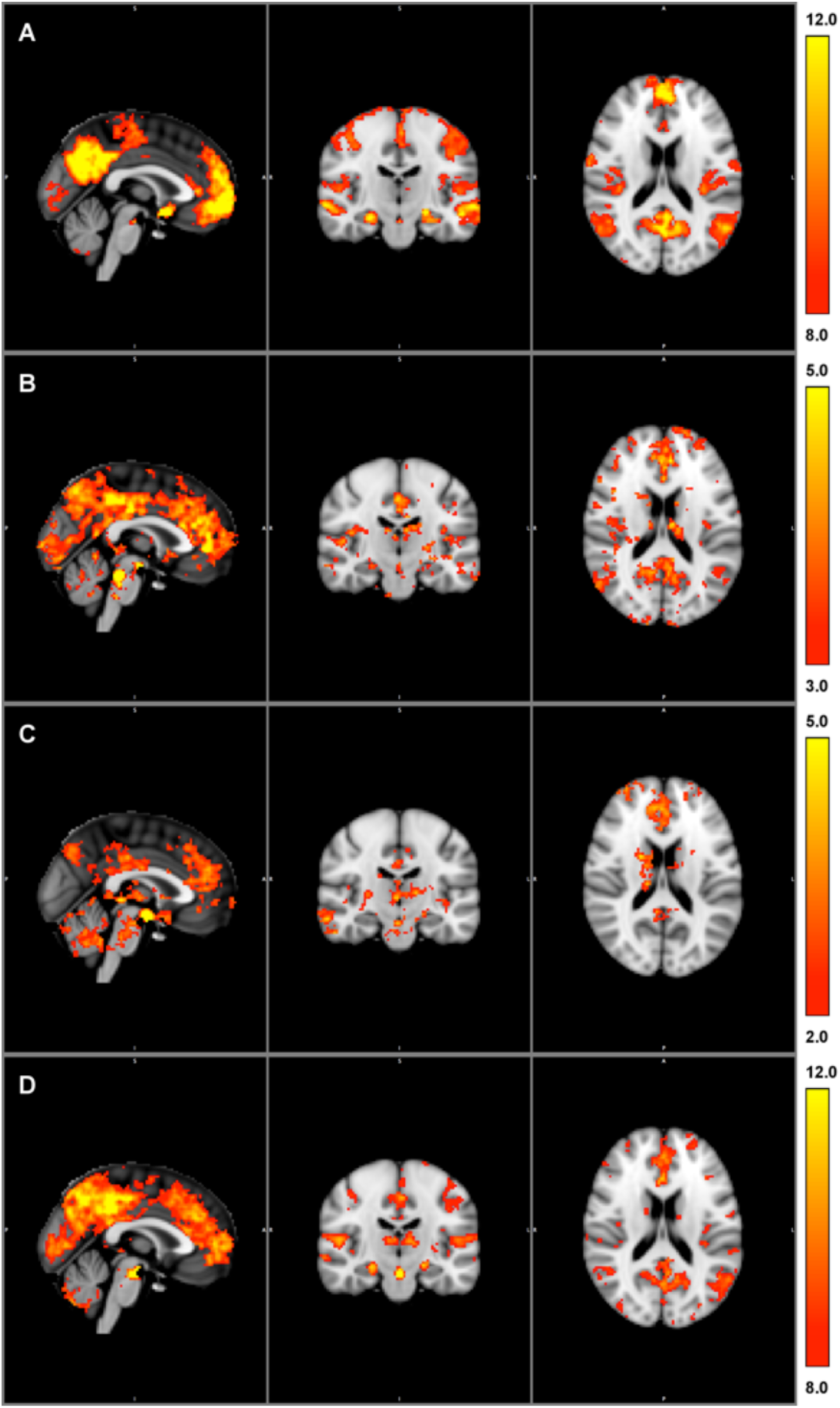
Seed based connectivity profiles using the MSDB (A), PnO (B), SuM (C) and VTA (D) as seeds. Slice positions are X=0, Y= −18 and Z = 18 mm in MNI coordinates. A Overlays are the t-values for significant voxels (p< 0.05) computed using fsl-randomise with threshold free cluster enhancement, however higher thresholds for the t-values are used in the display to show relevant details.

**Supplementary Figure 3.**
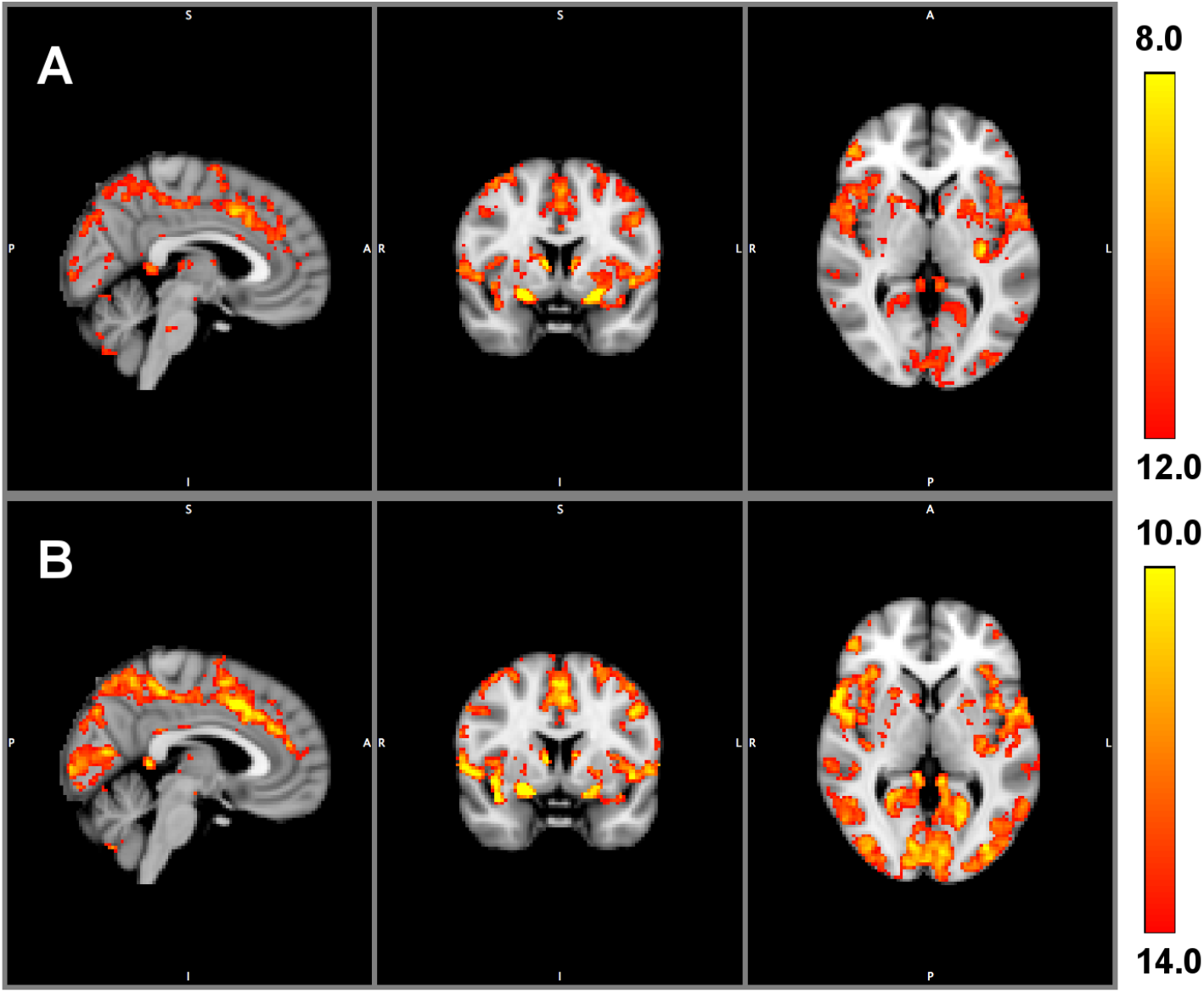
Seed based connectivity profiles using the Basal forebrain Left (A) and Right (B) as seeds. Overlays are the corrected t-values computed using randomise with threshold free cluster enhancement. All voxels displayed are significant at p < 0.05 using fsl-randomise permutation tests and threshold free cluster enhancement, however higher thresholds are used in the display to show relevant details. Slice positions are X=4, Y= 4 and Z = 2 mm in MNI coordinates.

**Supplementary Figure 4.**
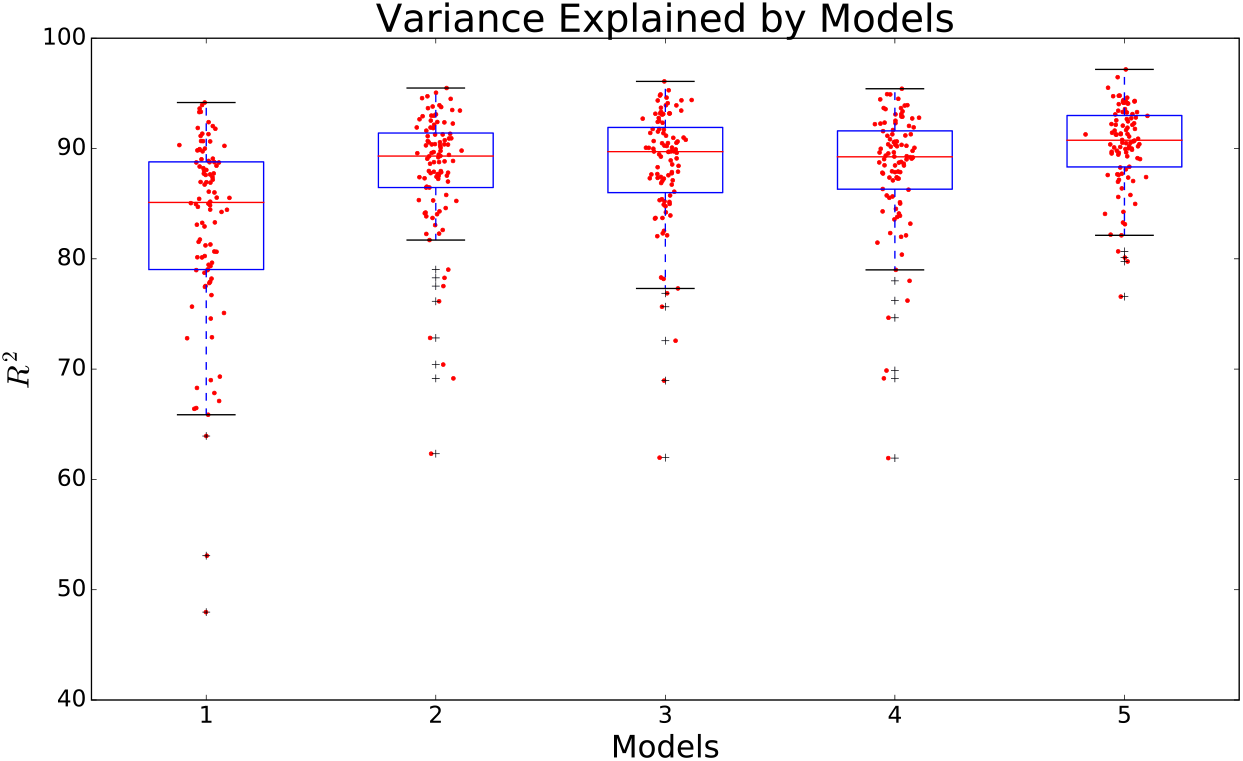
Boxplots of the explained variance for the DCM analyses. Model 1 is the basic DMN including PCC, aMPFC and pIPL. Model 2 includes basic DMN and MS + HC. Model 3 includes basic DMN, MS, HC, SuM and PnO. Model 4 includes basic DMN, MS, HC and PnO. Model 5 is the complete model.

**Supplementary Figure 5.**
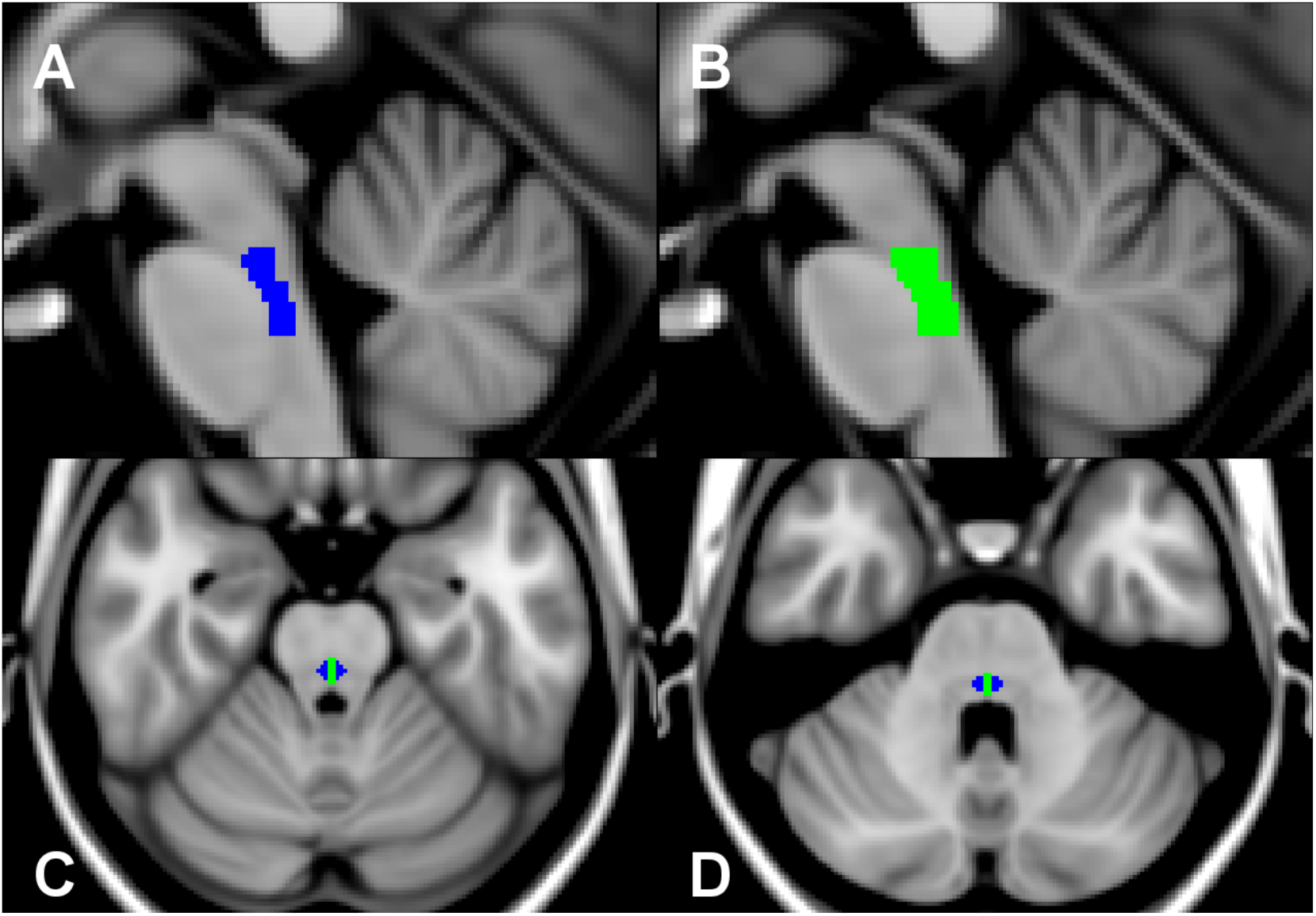
Serial sagittal sections ((A) x=2 (B) x=1) through the MNI single subject template showing the relative positions of the PnO (blue) and MRN (green) as delineated in the Harvard AAN atlas. Axial sections ((C) z=-32 and (D) z=−24) show the encapsulation of the MRN by PnO.

